# Development of a high-density 665 K SNP array for rainbow trout genome-wide genotyping

**DOI:** 10.1101/2022.04.17.488574

**Authors:** Maria Bernard, Audrey Dehaullon, Guangtu Gao, Katy Paul, Henri Lagarde, Mathieu Charles, Martin Prchal, Jeanne Danon, Lydia Jaffrelo, Charles Poncet, Pierre Patrice, Pierrick Haffray, Edwige Quillet, Mathilde Dupont-Nivet, Yniv Palti, Delphine Lallias, Florence Phocas

**Author notes:** **Correspondence:** Florence PHOCAS.

## Abstract

Single nucleotide polymorphism (SNP) arrays, also named « SNP chips », enable very large numbers of individuals to be genotyped at a targeted set of thousands of genome-wide identified markers. We used preexisting variant datasets from USDA, a French commercial line and 30X-coverage whole genome sequencing of INRAE isogenic lines to develop an Affymetrix 665 K SNP array (HD chip) for rainbow trout. In total, we identified 32,372,492 SNPs that were polymorphic in the USDA or INRAE databases. A subset of identified SNPs were selected for inclusion on the chip, prioritizing SNPs whose flanking sequence uniquely aligned to the Swanson reference genome, with homogenous repartition over the genome and the highest Minimum Allele Frequency in both USDA and French databases. Of the 664,531 SNPs which passed the Affymetrix quality filters and were manufactured on the HD chip, 65.3% and 60.9% passed filtering metrics and were polymorphic in two other distinct French commercial populations in which, respectively, 288 and 175 sampled fish were genotyped. Only 576,118 SNPs mapped uniquely on both Swanson and Arlee reference genomes, and 12,071 SNPs did not map at all on the Arlee reference genome. Among those 576,118 SNPs, 38,948 SNPs were kept from the commercially available medium-density 57K SNP chip. We demonstrate the utility of the HD chip by describing the high rates of linkage disequilibrium at 2 kb to 10 kb in the rainbow trout genome in comparison to the linkage disequilibrium observed at 50 kb to 100 kb which are usual distances between markers of the medium-density chip.

## 1 Introduction

Next-generation sequencing (NGS) has transformed the fields of quantitative, ecological and evolutionary genetics by enabling the discovery and cost-effective genotyping of thousands to millions of variants across the genome, allowing for genome-wide association studies (GWAS) of complex traits, genomic selection (GS) through accurate inference of relationships among individuals (Meuwissen and Goddard, 2010), inbreeding (Kardos et al., 2015), population structure and genetic diversity studies. Large numbers of densely genotyped individuals are required to get accurate results thanks to a high SNP density along the genome that constructs strong linkage disequilibrium between SNP and causative mutations (de Roos et al., 2008). However, regardless of the animal or plant species, it remains very challenging to cost-effectively genotype large numbers of individuals at polymorphic sites in all the genomes. An appealing strategy is to use a cheaper and reduced-density SNP chip with markers being chosen for optimizing the imputation accuracy to higher density genotypes. Genotype imputation describes the process of predicting genotypes that are not directly assayed in a sample of individuals (Marchini and Howie, 2010). Imputation has become a standard practice in research to increase genome coverage and improve GS accuracy and GWAS resolution, as a large number of samples can be genotyped at lower density (and lower cost) then imputed up to denser marker panels or to sequence level, using information from a limited reference population (Phocas, 2022).

Two main methods are employed for large-scale and genome-wide SNP genotyping. Array-based methods use flanking probe sequences to interrogate pre-identified SNPs (often named “SNP chips”). The alternative genotyping-by-sequencing (GBS) methods call SNPs directly from the genome (Davey et al., 2011). In GBS methods, either restriction enzymes are used to target sequencing resources on a limited number of cut sites (Baird et al., 2008) or low-coverage whole genome resequencing is performed. Low-coverage GBS followed by imputation has been proposed as a cost-effective genotyping approach for human genetics studies (Pasaniuc et al., 2012), as well as farmed species (Gorjanc et al., 2017) that cannot afford a high development of genomic tools. Nevertheless, compared to GBS methods, SNP chips offer a robust and easily replicable way of genotyping samples at a consistent set of SNPs, with very low rates of missing data.

Medium (~thousands to tens of thousands of loci) and high (~hundreds of thousands of loci) density SNP chips have been routinely developed for commercial species to perform genomic selection (Meuwissen et al., 2001) and to identify genes playing significant roles in livestock and crop performances (Goddard et al., 2016). SNP chips developed for model organisms or farmed species have also been utilised to address evolutionary and conservation questions, in particular in animal populations. For example, they have been used to identify signatures of adaptation in cattle (Gautier et al., 2010) or genes under selection in grey wolves (Schweizer et al., 2016), characterize the genetic diversity and inbreeding levels in pig (Silió et al., 2013), sheep (Mastrangelo et al., 2014), cattle (Rodríguez-Ramilo et al., 2015) or fish (D’Ambrosio et al., 2019), and infer the genomic basis of recombination rate variation in cattle (Sandor et al., 2012) or sheep (Johnston et al., 2016; Petit et al., 2017).

While there is now over ten fish and shellfish species for which commercial SNP arrays had been developed (Boudry et al., 2021), most of those contain only about 50 to 60K SNPs. Such medium-density chips are sufficient for genomic selection purposes but are clearly too low-density tools for fine QTL detection and help in identification of causal variants. As rainbow trout (*Oncorhynchus mykiss*) is a major academic model for a wide range of investigations in disciplines such as cancer research, toxicology, immunology, physiology, nutrition, developmental or evolutionary biology in addition to quantitative genetics and breeding (Thorgaard et al., 2002), it is important to get access to very high-density genomic tools for this salmonid species.

For rainbow trout, SNP discovery has been firstly done through sequencing of restriction-site associated DNA (RAD) libraries (Palti et al., 2014), reduced representation libraries (RRL) (Sánchez et al., 2009) and RNA sequencing (Sánchez et al., 2011). A first commercial medium-density Axiom® Trout Genotyping array (hereafter termed 57K chip) has then been developed (Palti et al., 2015) and produced by Affymetrix (Thermofisher). Since then it has been largely used in population genetics studies (Larson et al., 2018; D’Ambrosio et al., 2019; Paul et al., 2021), GWAS and GS accuracy works for various traits in farmed populations (Gonzalez-Pena et al., 2016; Vallejo et al., 2017, 2019; Reis Neto et al., 2019; Rodríguez et al., 2019; Yoshida et al., 2019; Fraslin et al., 2019, 2020; Karami et al., 2020; D’Ambrosio et al., 2020; Blay et al., 2021a, 2021b). However, out of the 57,501 SNPs included in this chip, nearly 20,000 were found to be unusable because they were either duplicated due to the ancestral genome duplication or showing primer polymorphism in 5 French commercial or experimental lines (D’Ambrosio et al., 2019). Of the 57,501 markers from the original chip, 50,820 are uniquely localized on the Swanson reference genome (Pearse et al., 2019), and in the remaining number, only 38,332 markers pass the control quality filters (no primer polymorphism, call rate> 97%, Minor Allele Frequency (MAF)> 0.001 over 3,000 fish from 5 French lines).

To overcome these limitations as well as to get access to a more powerful tool for GWAS and population genetics studies in rainbow trout, the aim of our study was to develop a high-density SNP array. To develop this resource for rainbow trout, we made use of a large set of resequencing data from 31 doubled haploid (DH) lines from Washington State University (WSU) and Institut National de la Recherche pour l’Agriculture, l’Alimentation et l’Environnement (INRAE). In the USA, 12 WSU DH lines have been created by androgenesis (Young et al., 1996) while in France 19 DH INRAE lines (called isogenic lines) were produced by gynogenesis (Quillet et al., 2007). The 12 WSU DH lines as well as 7 of the INRAE isogenic lines served as basic material for the variant search and SNP selection for the 57K chip (Palti et al., 2015).

In this study, we describe how we overcame the limitations of duplications in the rainbow trout genome, in order to identify and locate polymorphisms. We describe the subset of detected SNPs that was selected for inclusion on a custom high-density SNP chip. It was used to genotype 463 samples from two different French commercial populations. We test the genotyping success rates, that is, the proportion of SNPs included on the array that are polymorphic and successfully genotyped. We demonstrate the utility of this SNP chip to infer linkage disequilibrium in the genome of this species.

## 2 Materials and methods

### 2.1 Use of the USDA database for initial SNP detection

Gao et al. (2018) constituted a first large SNP database (USDA1) by performing high coverage whole genome resequencing (WGS) with 61 unrelated samples, representing a wide range of rainbow trout and steelhead populations. Of the 61 samples, 11 were doubled-haploid lines from Washington State University (WSU), 12 were aquaculture samples from AquaGen (Norway), 38 were from wild and hatchery populations from a wide range of geographic distribution (Califormia, Oregon, Washington and Idaho states in the USA; Canada; Kamtchatka Penninsula in Russia). Overall, 31,441,105 SNPs were identified with 30,302,087 SNPs located on one of the 29 chromosomes of the Swanson reference genome assembly (Omyk_1.0; GenBank, assembly accession GCA_002163495.1) (Pearse et al., 2019).

A second database (USDA2) with 17,889,078 SNPs coming from resequencing of 24 USDA samples was added to the initial USDA1 database. The samples were composed of 12 representatives from the USDA-NCCCWA odd-year class and 12 from the even-year class as previously described (NCBI BioProject PRJNA681179; Liu et al., 2021). The SNP discovery analysis followed the methods of (Gao et al., 2018).

By merging these two databases using BCFtools (Danecek et al., 2021), we constituted a single USDA database that contained 35,732,342 distinct SNPs, with 34,170,401 placed on the 29 chromosomes or mitochondrial chromosome of the Swanson reference genome. SNP filtering was performed to remove non bi-allelic variants and SNPs with MAF < 1% using a homemade python script. The final USDA clean database contained 29,024,315 SNPs.

### 2.2 Whole genome resequencing of INRAE isogenic lines and use of the INRAE database for SNP detection

Genomic DNA was extracted from fin clips of 19 rainbow trout INRAE isogenic lines. Whole-genome paired-end sequencing libraries were prepared and sequenced using the Illumina HiSeq 2000, Hi Seq 3000 or HiSeq X-Ten platforms at a depth of genome coverage ranging from 10X to 32X per sample. The 19 isogenic lines were sequenced in two batches that were processed successively. The first batch contained sequencing data from 12 samples (doubled haploid individuals) coming from 11 isogenic lines. The second batch contained sequencing data from 17 samples (doubled haploid individuals) coming from 17 isogenic lines (9 lines already sequenced in batch 1; and 8 lines not previously sequenced). Overall, 10 out of the 19 isogenic lines were sequenced twice. This resulted in a total of 8,911,630,867 paired reads with a median of 321,575,464 per sample.

Sequence reads from each of the 12 samples from the first batch were mapped to the Swanson rainbow trout reference genome (GenBank assembly accession GCA_002163495.1; Pearse et al., 2019) using BWA MEM v.0.7.12 (Li, 2013). We then ran Samtools sort (v1.3.1, (Danecek et al., 2021)) to sort the alignment data by chromosome and scaffold locations. Afterwards, PCR duplicates were marked using Picard Tools (v.2.1.1, Broad Institute 2019) MarkDuplicates. Variant calling was then performed for each sample using GATK (v3.7; McKenna et al., 2010) HaplotypeCaller (options *-stand_call_conf 30 -mbq 10*), leading to 12 vcf files. A variant reference file containing 1,207,861 high quality SNPs was generated by keeping variants with *QUAL>=1050* from the vcf files. This file was then used for the recalibration step, using GATK BaseRecalibrator and PrintReads. The recalibrated BAM files were then used as input for the variant calling step using GATK HaplotypeCaller in ERC GVCF mode. The resulting 12 GVCF files were then merged into a single vcf file containing 24,944,575 variants using GATK GenotypeGVCFs. The vcf file was then filtered as follows using GATK VariantFiltration: *DP<120; MQ<30.0; QUAL<600; AN<12*. To filter out putative PSVs (Paralogous Sequence Variants), we filtered out variants with heterozygous genotypes in at least two of the 12 doubled haploid samples. The filtered vcf file from the first batch contained 11,113,836 variants.

The second samples sequence batch were analyzed following the same procedure as for the first batch with few updates. Prior to sequence alignment, sequences have been filtered using trimmomatic 0.36 (Bolger et al., 2014) to remove Illumina Truseq adapters, trim low quality bases, keep trimmed reads with a sufficient length and average quality. These parameters removed 3.8% of the reads, keeping 6,349,173,142 reads over the 17 samples. Alignment software was updated to use BWA MEM v.0.7.15. First calling to create a high quality variants set to recalibrate the BAM files was avoided by directly using the final vcf file from the first batch analysis. These recalibrated BAM have been submitted to GATK Haplotype caller as before to generate GVCF files. To increase confidence in the SNP calling, we also added 2 other SNP callers: Samtools mpileup and FreeBayes 1.1.0 (Garrison and Marth, 2012). GATK calling results were jointly genotyped using GATK GenotypeGVCFs on the 12 GVCF files from the first batch and the 17 newly generated GVCF files. This calling procedure resulted in 3 VCF files, one for each caller. Calling from GATK contained 29 samples (from the 19 isogenic lines, i.e. with 10 lines replicated) and 31,454,943 variants; Freebayes and Mpileup was used only on the second batch and contained 19 samples and 25,805,271 and 30,340,281 variants respectively.

The final step for variant calling was to intersect the 3 calling datasets using VCFtools_0.1.12a (Danecek et al., 2011), to keep only variants called by the 3 callers (genotypes kept were the GATK ones). SNP and INDEL were separated using GATK Select Variants, and SNP were filtered with GATK VariantFiltration by following the GATK recommendations (*QD < 2.0 || MQ < 40.0 || FS > 60.0 || SOR > 3.0 || MQRankSum < −12.5 || ReadPosRankSum < −8.0*). This constitutes the INRAE1 variants dataset which includes 14,439,713 SNPs.

Using a homemade python script to parse VCF file, INRAE1 dataset was filtered to keep only bi-allelic SNP localized on the 29 trout chromosomes or mitochondrial chromosome, fully genotyped for all 29 samples. As 10 isogenic lines were duplicated, we also checked genotype consistency and removed SNP with more than 1 isogenic line genotype discordance. Finally, we kept one sample per isogenic line (with the deepest sequencing) and filtered out SNP with more than 1 heterozygote genotype as they may represent duplicated genome regions. Among the 14,439,713 variants, we kept 10,286,009 SNPs (71.23%).

We merged them using BCFtools with a second dataset INRAE2, containing 14,478,077 SNPs called from 60 samples of a commercial line from ”Les Fils de Charles Murgat” (Beaurepaire, France) and whose resequencing was described in Fraslin et al. (2020).

This merged dataset was filtered like the merged USDA dataset, to keep bi-allelic SNP localized on the 29 trout chromosomes of the Swanson reference genome or mitochondrial chromosome, with a MAF > 1%. The final INRAE cleaned database contained 16,466,188 SNPs.

### 2.3 Merging the USDA and INRAE SNP databases and SNP preselection

A total of 32,372,492 distinct SNPs were selected for consideration for the HD chip, from a combination (BCFtools merge) of the USDA and INRAE databases (Supplementary data 1).

An overview of the process to detect and select SNPs for inclusion on the array is provided in Figure 1.

**Figure 1.**
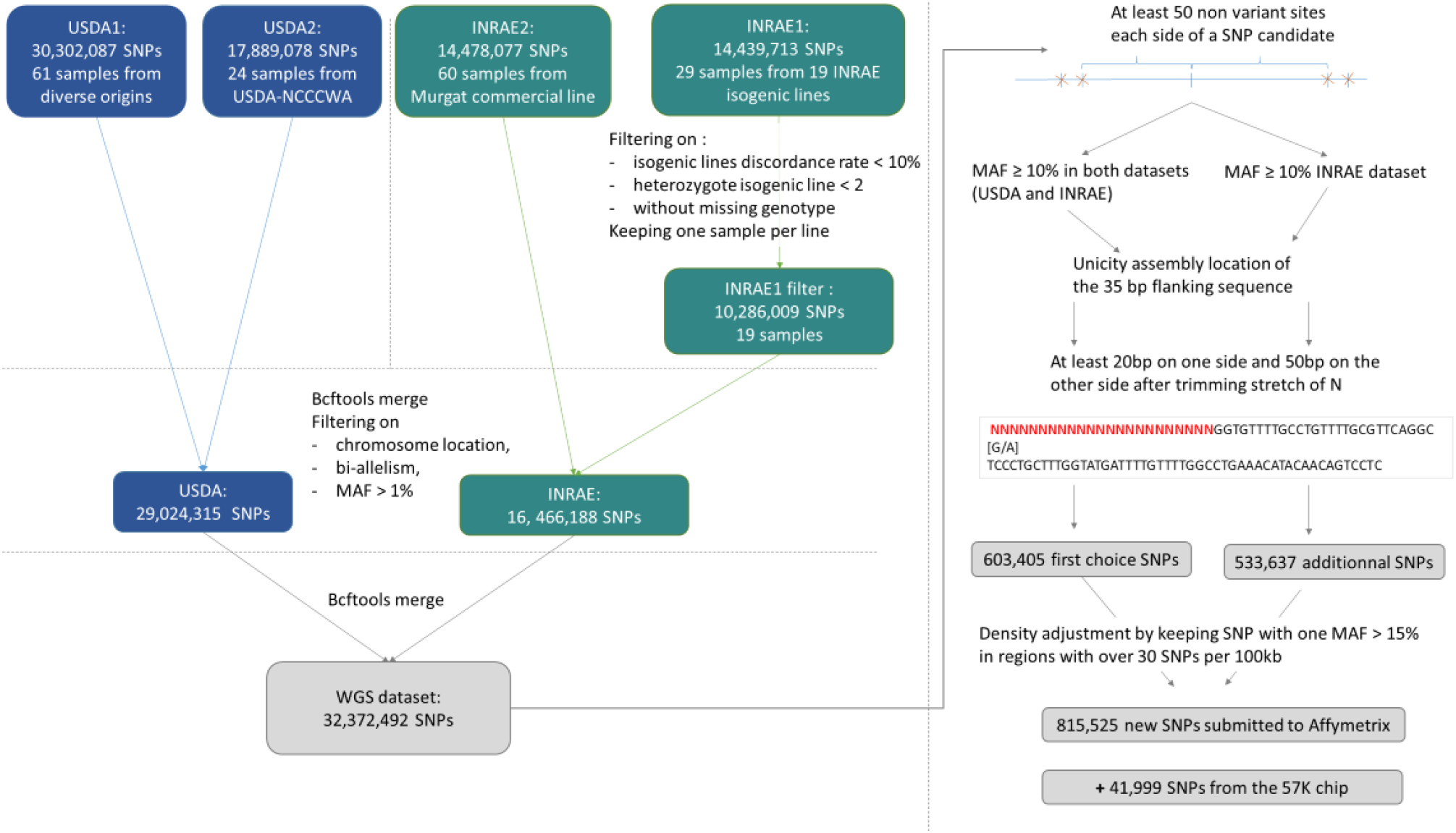
Process for submitted SNPs for inclusion on a high-density genotyping array

SNPs were further filtered to be at least 50 base pairs from the closest identified SNP which resulted in a subset of 3,679,547 SNPs.

During vcf files merging, additional alleles may be added on shared variant positions and some variants previously removed from the INRAE dataset (on replicate discordance or high isogenic heterozygosity rate) may be reincluded. Thus, for a first SNP preselection, in addition to filtering SNPs with MAF ≥10% in both the USDA and INRAE databases, we applied filters on bi-allelism variant and on a maximum number of 4 heterozygote INRAE isogenic lines.

Assessment included a check for duplicate flanking information suggesting repetitive elements, and an assessment of the complexity of the flanking sequence:

i. unicity of at least one side 35 bp-sequence for each SNP. This was done by blasting (Camacho et al., 2009) the 35 bp on the reference assembly genome and by checking that the best match was unique and located on the expected chromosome.
ii. trimming of each side 50 bp sequence if it contained more than 3 successive N. Variants were kept if at least the shortest trimmed sequence contained 20 bp and the other 50 bp (homemade python script).

This first high-quality selection represented 633,405 SNPs. Trimmed flanking sequence each side of the SNP was extracted for all SNPs and formatted for Affymetrix (Thermo Fisher Scientific, USA) according to their specifications.

From this first submission to Affymetrix quality control, only 457,086 SNPs were qualified as recommended to be designable for the HD array and among them, only 351,755 were not ambiguous, meaning they were not of the type [G/C] or [T/A] that would require 4 probes instead of only two to distinguish the alleles.

To get sufficient recommended variants and to avoid the selection of markers that will use twice the space used by the others on the HD array, we decided to resubmit a large second set of variants to Affymetrix quality check. The same procedure was applied to produce a second more relaxed preselected set of SNPs by keeping SNPs with a MAF ≥ 10% in the INRAE dataset only. This second preselection contained 533,637 additional SNPs. Among that additional set, 134,086 SNPs were specific to the INRAE dataset while the others were also present in the USDA dataset but with MAF below 10%.

We merged the first recommended set of 457,086 SNPs with this additional set of 533,637 SNPs. Then we removed all ambiguous SNPs of type [G/C], [C/G], [T/A] or [A/T]. Finally, densities were adjusted such that in regions with more than 30 SNPs retained per 100 kb by the previous filters, we only kept SNPs with MAF ≥15% in at least one of the two INRAE or USDA databases.

This procedure resulted in a selection of 815,525 SNPs for the final submission in October 2020 to Affymetrix for assessment of the suitability of the SNPs for inclusion on a custom AXIOM 96HT SNP chip. Of the submitted SNPs, a total of 623,544 SNPs were deemed to be “designable” (recommended or neutral) in either the forward or reverse flanking sequence based on the Affymetrix pconvert score.

### 2.4 Keeping informative variants from the medium-density Axiom® Trout Genotyping array

The INRAE and USDA research teams were willing to keep in the HD chip design the informative markers from the 57K chip. Therefore 41,999 SNPs out of its 57,501 SNPs were designable in either forward and reverse directions and were kept for the HD chip design (Supplementary Data 2).

At the only exception of 8 specific SNPs, all the markers had a unique position on the Swanson reference genome and MAF > 5% in at least one French or North American population. Among them, 38,826 SNPs were also put on a 200K chip that was built on 120 resequenced mostly “wild” genomes from over 40 locations from Russia, Alaska Canada down through Washington, Oregon and California (Ben Koop’s personal communication).

### 2.5 Selection of SNPs for the HD-trout SNP chip

In total 664,531 SNPs corresponding to 701,602 probesets (some SNPs were tiled in both directions as both their forward and reverse flanking sequence was assessed to be neutral) passed the Affymetrix final quality control to be designed on the custom HD Axiom array. Only 40,987 of the 41,999 SNPs from the 57K chip remained on the HD final design.

Among the selected SNPs, 664,503 were mapped on the 29 chromosomes of the Swanson reference genome (Figure 2), while 28 were positioned on the mitochondrial genome.

**Figure 2.**
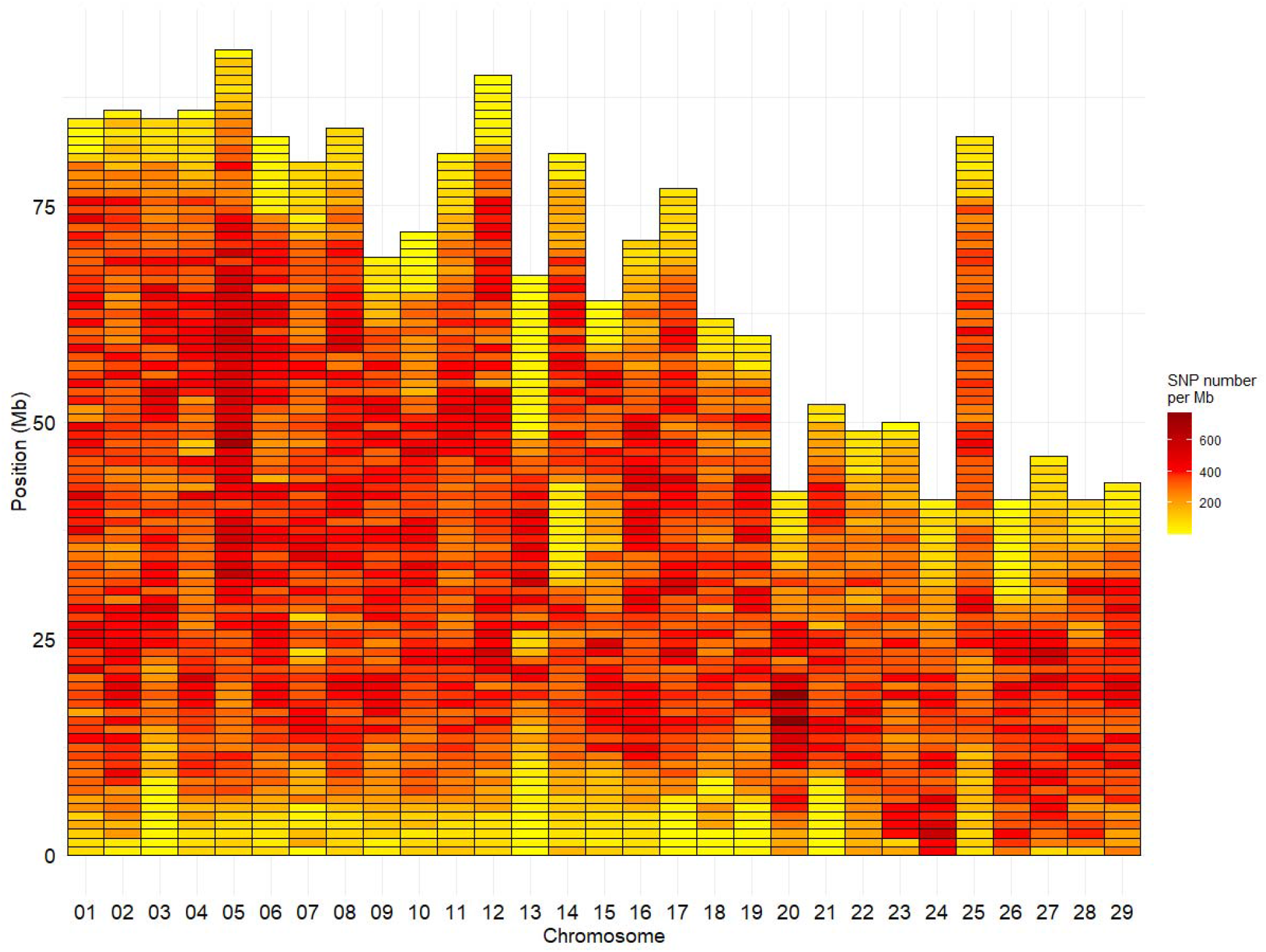
Marker density per Mb for the HD Trout Affymetrix array with 664,503 SNPs positioned on the 29 chromosomes of the Swanson genome reference

Based on the Swanson reference genome mapping, the average (median) SNP density on the chromosomes was 293 (324) SNPs per Mb (Figure 2), with SNP density varying from 2 to 774 SNPs per Mb. The average (median) intermarker distance was 2.9 kb (1.3 kb). Maximum intermarker distance was 243 kb and only 5.4% of intermarker distance was over 10 kb (0.02% over 100 kb).

### 2.6 Samples for genotyping

This study used fin samples collected from “Bretagne Truite” (Plouigneau, France) and “Viviers de Sarrance” (Sarrance, France) commercial lines, hereafter named LB and LC lines respectively, that were sampled for the FEAMP project Hypotemp (n° P FEA470019FA1000016).

Pieces of caudal fin sampled from 463 fish (288 from LB line and 175 from LC line) were sent to Gentyane genotyping platform (INRAE, Clermont-Ferrand, France) for DNA extraction using the DNAdvance kit from Beckman Coulter following manufacturer instructions and genotyping using the newly constructed HD SNP array.

The first round of quality control was done by ThermoFisher software AxiomAnalysisSuite™ with threshold values of 97% for SNP call rate and 90% for sample call rate. All the 288 individuals of the LB line passed the preliminary control, while 174 out of the 175 individuals from LC line passed the control quality.

Following array hybridization and imaging, genotypes were called using default settings in the Axiom Analysis Suite software and exported from the software in PLINK (Purcell et al., 2007) format. The 701,602 SNP probe flanking sequences were realigned to the new Arlee reference genome using BLAST. Indeed, recently USDA/ARS (Gao et al., 2021) released a second reference genome assembly (GCA_013265735.3) for *Oncorhynchus mykiss* as long reads-based de-novo assembly for a second WSU DH line, named Arlee line, had been performed. Because Arlee lineage was closer than Swanson lineage from the INRAE isogenic lines (Palti et al., 2014), it was decided to keep for further analysis only the SNPs that were mapped uniquely on one of the 32 chromosomes of this new reference genome.

In addition, we used the WGS information of 20 samples sequenced in Gao et al. (2018)’ study (with average genome coverage above 20X) to extract their genotypes for SNPs included in the HD chip and positioned on the Arlee reference genome (Gao et al., 2021; USDA_OmykA_1.1; GenBank, assembly accession GCA_013265735.3). Those samples came from hatchery (Dworhak, L. Quinault, Quinault, Shamania) and wild (Elwha) populations from the North-West of USA (4 samples for each of the 5 populations) and were proved to be genetically close to each other and very distant from the Norwegian Aquagen aquaculture population (Gao et al., 2018). The idea was to infer and compare the level of linkage disequilibrium across the HD markers from wild/hatchery American populations and farmed French selected lines.

### 2.7 Allele frequencies and linkage disequilibrium across populations

We then used PLINK v1.9 (www.cog-genomics.org/plink2) to calculate allele frequencies, filter SNPs at low MAF or individuals with high identity by descent (IBD) values and derive linkage disequilibrium (LD) measured as the correlation coefficient r2, using the mapping of the SNP probe flanking sequences to the Arlee genome.

Allele frequencies were calculated per population for each SNP. SNPs were then filtered to only those with a MAF ≥ 5%, leaving 249,055 variants for American populations, 420,778 SNPs for the LB line, and 423,061 SNPs for the LC line. The set of individuals was also filtered using *“rel-cutoff 0.12”* to exclude one member of each pair of samples with observed genomic relatedness above 0.12, keeping 120 samples across populations, corresponding to 20, 45 and 55 individuals for American populations, LB and LC lines, respectively. Linkage disequilibrium (r2) between all pairs of SNPs on the same chromosome and at physical distances up to 1 Mb was then calculated using the PLINK options *‘--r2 --ld-window 50000 –ld-window-kb 1001 –ld-window-r2 0.0’*. The r2 values were binned into 2 kb units and per-bin averages calculated using R (R Core Team, 2019) for all chromosomes. The LD decay over physical distance up to 100 kb was then plotted in R.

## 3 Results

### 3.1 SNP identification and characterization on the joint USDA+INRAE WGS database based on the Swanson reference genome

Density of SNPs varied strongly from one chromosome to another with average SNP density per Mb ranging from 13,200 for Omy26 to 20,132 for Omy22. Across all chromosomes, the average SNP density per Mb was 16,483 SNPs (Figure 3). The Mb with the minimum density contained 451 SNPs while the Mb with the highest density contained 31,819 SNPs.

**Figure 3.**
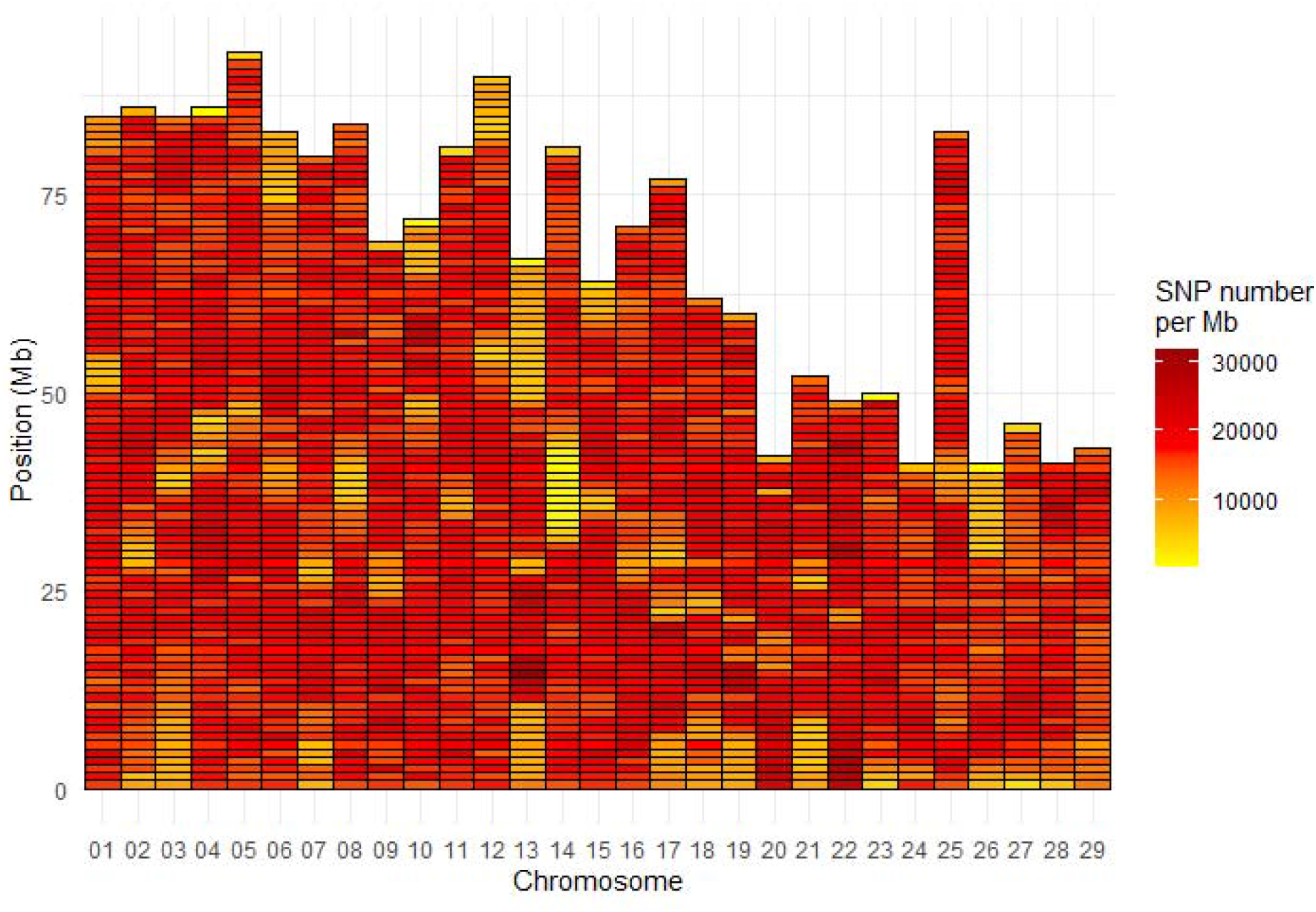
SNP density per Mb for the INRAE_USDA full variant dataset (32.4M SNPs) located on the 29 chromosomes of the Swanson genome reference

SNP identified in USDA or INRAE databases differed in terms of MAF distribution (Figure 4): 70% and 49% of SNPs had a MAF below 15% (40% and 15% had a MAF below 5%, respectively) while only 9.5% and 18% of SNPs had a MAF above 35% in the USDA and INRAE datasets respectively.

**Figure 4.**
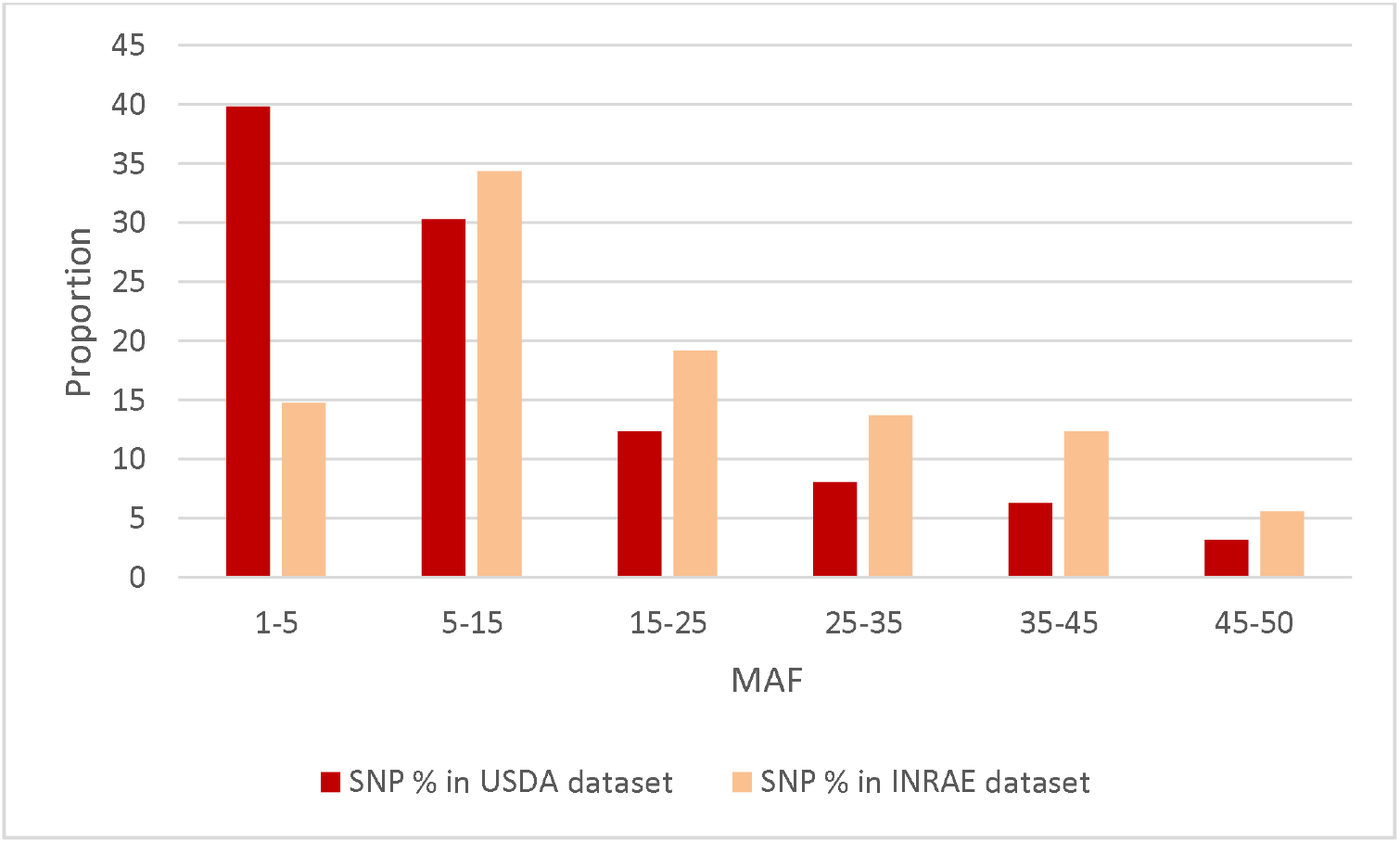
MAF distribution of USDA or INRAE SNP datasets. These datasets have been filtered to keep bi-allelic SNP with a minimal MAF > 1% in their respective populations.

### 3.2 HD chip

Based on genotyping the 288 LB samples, 65.34% of markers were polymorphic, had individuals with all three genotypes, and passed Affymetrix filtering metrics in the Axiom Analysis Suite software to be categorized as “PolyHigh Resolution” variants. Of those that “failed” to be in that category, 15.35% passed filtering metrics but were monomorphic, 10.71 % passed filtering metrics but the minor allele homozygote was missing, and the remainder 8.60% failed due to low call rates or other quality filters. The total number of best recommended markers was 91.81% corresponding to 610,115 SNPs out of the 664,531 genotyped variants.

Based on genotyping the 175 LC samples, 69.91% of markers were polymorphic, had individuals with all three genotypes, and passed Affymetrix filtering metrics in the Axiom Analysis Suite software to be categorized as “PolyHigh Resolution” variants. Of those that “failed” to be in that category, 5.63% passed filtering metrics but were monomorphic, 14.84 % passed filtering metrics but the minor allele homozygote was missing, and the remainder 9.62% failed due to low call rates or other quality filters. The total number of best recommended markers was 90.86% corresponding to 603,768 SNPs out of the 664,531 genotyped variants.

Of the 664,531 SNPs which passed the Affymetrix quality filters and were included on the HD chip, 576,118 SNPs mapped uniquely on both reference genomes, and 12,071 SNPs did not map at all on the Arlee reference genome. Supplementary data 3 indicates both positions on the Swanson and Arlee reference genomes. Among those 576,118 SNPs, 38,948 SNPs were kept from the initial 57K chip.

On the Arlee mapping (GCA_013265735.3), the average SNP density on the chromosomes was one SNP per 3.8 kb, or 266 SNPs per Mb. The median intermarker distance was 1.5 kb with only 7% of the distances between successive markers being above 10 kb. The largest gap was 4.16 Mb at the end of chromosome Omy6, the second largest gap was 2.94 Mb at the end of chromosome Omy10 and the third largest gap was 2.75 Mb at the end of chromosome Omy13 (Figure 5). Only five other gaps were above 2 Mb with values ranging from 2.3 to 2.5 Mb on chromosomes Omy7, Omy10, Omy15 and Omy21.

**Figure 5.**
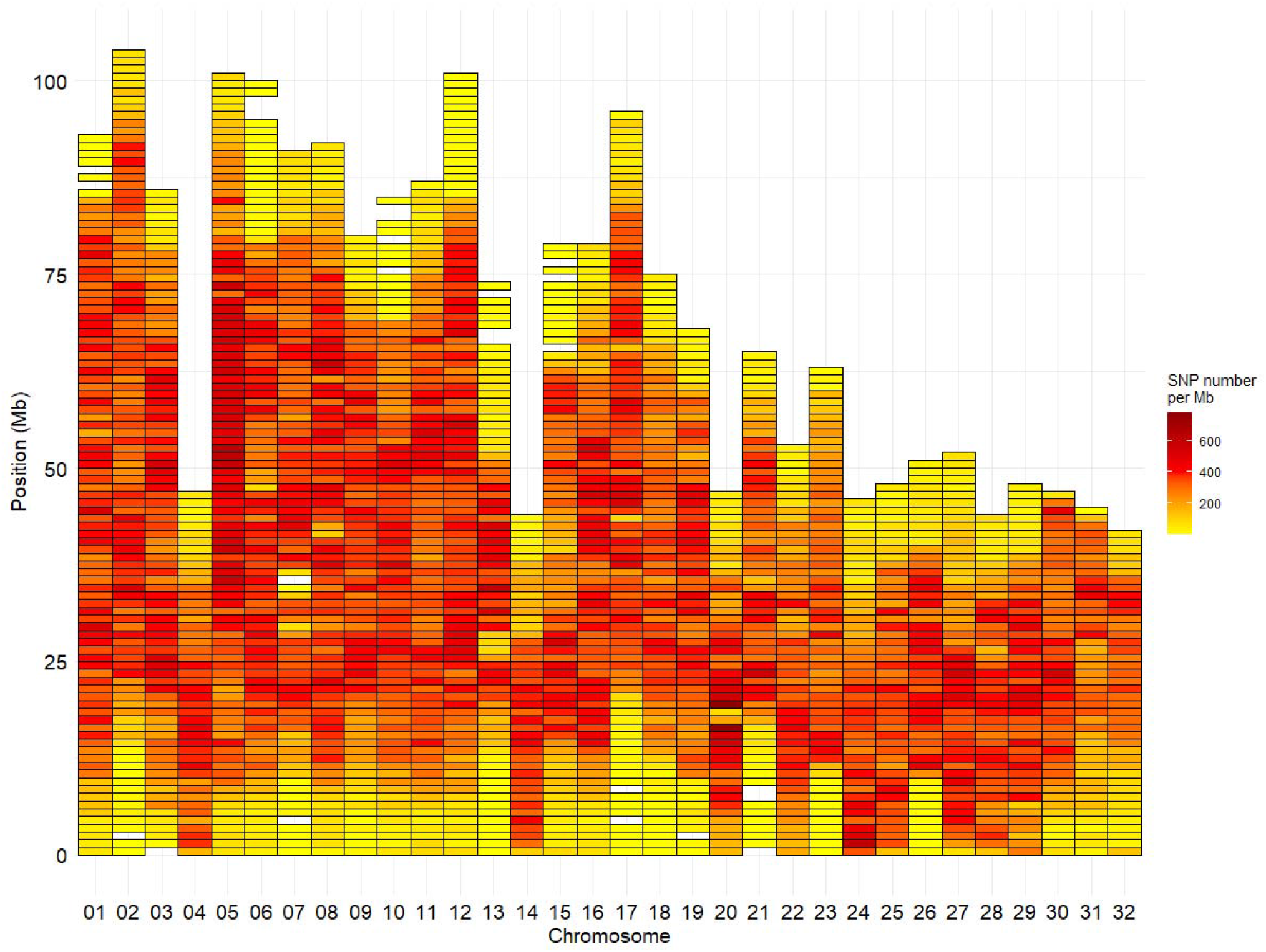
Marker density per Mb for the HD Trout Affymetrix array with 576,118 SNPs positioned on the 32 chromosomes of the Arlee reference genome

Finally, PLINK v1.9 software (www.cog-genomics.org/plink2) was used for a final SNP filtering based on keeping for further analysis SNPs with call rate above 95% and a deviation test from Hardy-Weinberg equilibrium (HWE) with a p-value < 10e-7 within each population. For LB line, 571,319 markers (474,937 being polymorphic) were kept after removing 1,136 miss genotyped SNPs and 3,663 ones with severe deviation from HWE. For LC line, 569,030 markers (487,940 being polymorphic) remained after removing 2,574 miss genotyped SNPs and 4,592 ones with severe deviation from HWE.

Regarding the American sequenced population, we extracted from the vcf files the genotypes for the 576,118 SNPs that were retained on the HD chip. Only 338,660 of those markers were polymorphic in the American population.

### 3.3 MAF distribution in the two French HD genotyped populations

Compared to variants called from sequence data, the MAF distribution of the HD selected SNPs was skewed to common alleles (Figure 6) with over 70% of SNPs with MAF above 5% in each of the two populations, and over 20% of SNPs with MAF over 35% in both populations. Among polymorphic SNPs (MAF > 0.001), the average (median) MAF was 24.1% (23.6%) in the LB line and 23.0% (21.5%) in the LC line.

**Figure 6.**
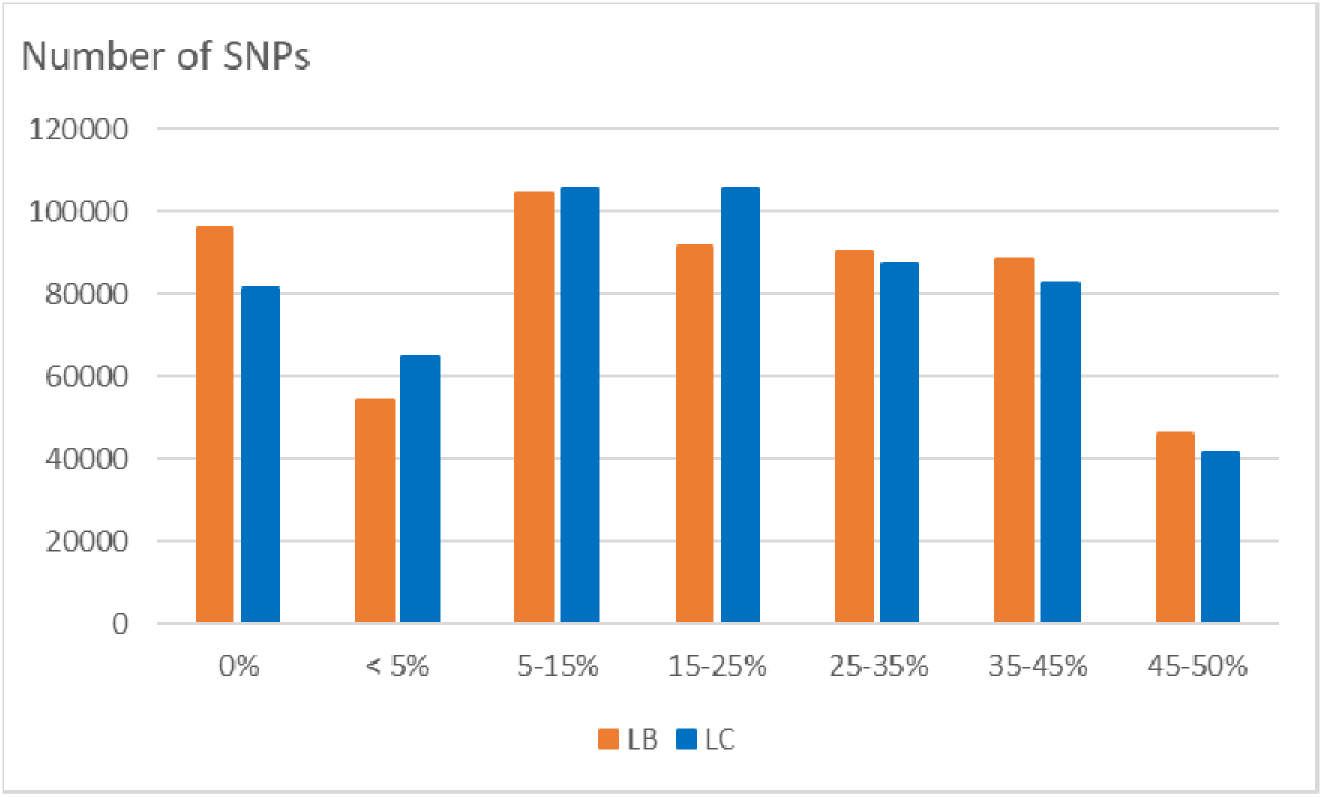
Distribution of SNPs according to their MAF class in the LB and LC French commercial lines.

### 3.4 Linkage disequilibrium analysis

The median intermarker distance was 2kb and the corresponding average r2 between neighbouring markers was 0.47, 0.44, and 0.36 in LB, LC and American population, respectively. As expected, average r2 tended to decrease with increasing distance between pairs of markers in all populations studied, the most rapid decline being over the first 10 kb (Figure 7). Linkage disequilibrium was very high, with r2 reaching 0.42, 0.39, and 0.27 at the average intermarker distance (4kb) for LB, LC, and American population respectively; at 50 kb distance, r2 average values were 0.32, 0.29, and 0.14 (Figure 7). At 500 kb, values were 0.25, 0.21, and 0.18 and values were still 0.22, 0.19, and 0.11 at 1 Mb, respectively for LB, LC and the American population (Figure 8). However, those r2 values may vary strongly from one chromosome to another as shown on Figure 8 for chromosomes Omy5 and Omy13 with respectively higher and lower linkage disequilibrium observed in comparison to the average values derived for all chromosomes.

**Figure 7.**
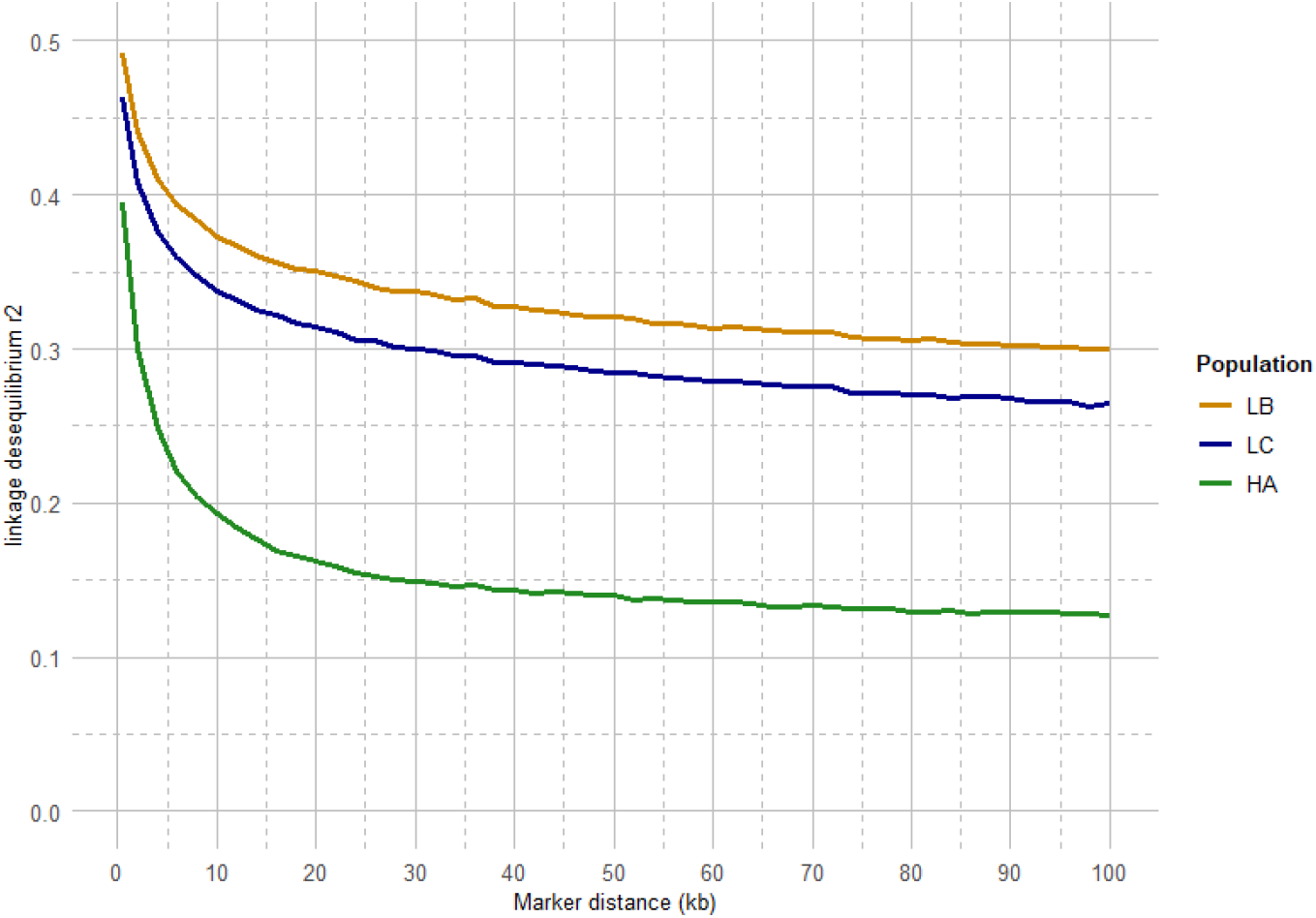
LD decay from 2 to 100 kb intermarker distances (average over the 32 chromosomes) for the LB and LC French commercial lines and the HA American population.

**Figure 8.**
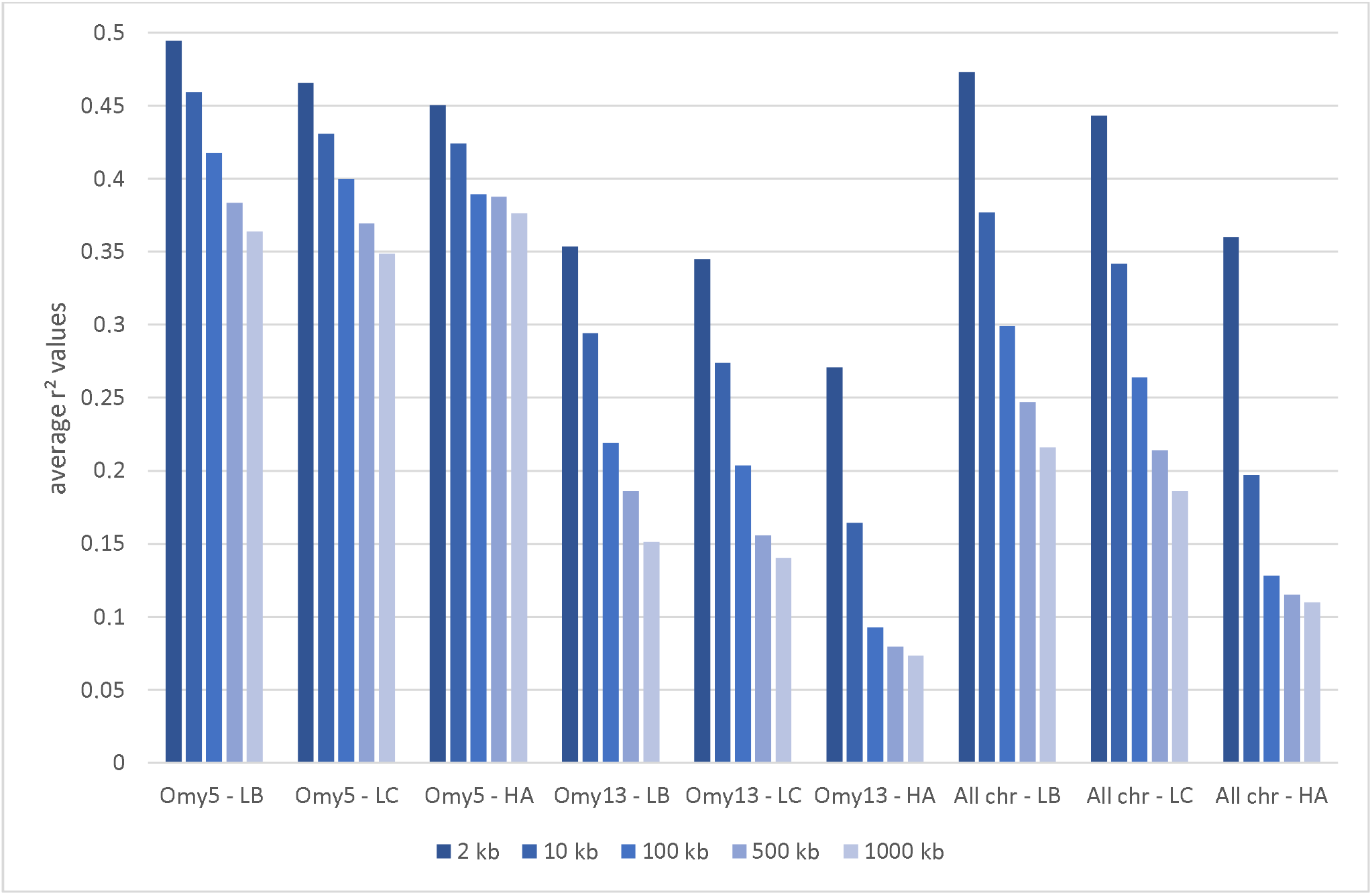
Average linkage disequilibrium (r2 values) from 2 to 1,000 kb derived for all chromosomes and only for Omy5 or Omy13 in populations LB, LC and HA, respectively

## 4 Discussion

In this study, based on the resequencing of tens of individuals from a diverse range of populations, we developed a high density (665K) SNP array that will be used for numerous applications, including genomic populations studies, GWAS or genomic selection. In fish, the first very high-density chip, named 930K XHD Ssal array, was developed for Atlantic Salmon using 29 fish from Aquagen lines and was a powerful tool to identify the key role of *VGLL3* gene on age at maturity (Barson et al., 2015) or the epithelial cadherin gene as the major determinant of the resistance of Atlantic salmon to IPNV (Moen et al., 2015). A similar approach was used in Atlantic salmon with whole genome re-sequencing of 20 fish from three diverse origins to generate a catalogue of 9.7M SNPs that were then filtered to design a 200K SNP chip (Yáñez et al., 2016). A similar number of 9.6M SNPs were identified for the development of a 700K SNP chip in catfish (Zeng et al., 2017). Recently, a set of 82 fish were collected from six different locations of China and re-sequenced to identify 9.3M SNPs to design a 600K SNP chip for large yellow croaker (Zhou et al., 2020).

Based on the resequencing of 85 samples by USDA and 79 samples by INRAE, we identified 32,372,492 SNPs that were variants (MAF ≥1%) in either the USDA or the INRAE sets. More precisely, 29.0 and 16.4 million SNPs were identified in the USDA and INRAE datasets respectively for equivalent number of sequenced individuals. The higher number of SNPs detected in the USDA dataset probably resulted from the larger number of diverse populations included in the USDA dataset. The USDA database included 11 doubled haploid individuals and 50 individuals from 7 commercial, hatchery or wild populations, compared to the INRAE database that included 19 doubled haploid individuals derived from one experimental line and 60 individuals sampled from a single French commercial line. For comparison purposes, the influence of the numbers of sequenced individuals and populations or breeds on the number of identified SNPs can be exemplified in two large-scale projects, the 1000 human genomes project and the 1000 bull genomes project. In the human genome, a pilot phase identified ~15 million SNP based on the WGS of 179 individuals from four populations (The 1000 Genomes Project Consortium, 2010); increasing the number of sequences to 2,504 coming from 26 populations across the world increased considerably the number of identified SNPs to 84.7 million (The 1000 Genomes Project Consortium, 2015; Fairley et al., 2020). Similarly, the first phase of the 1000 bull genomes project identified 26.7 million SNPs based on the resequencing of 234 bulls from 3 breeds (Daetwyler et al., 2014); again, the number of SNPs increased to 84 million by sequencing 2,703 individuals from 121 breeds (Hayes and Daetwyler, 2019). Another study in chicken highlights the importance of sequencing a diverse set of individuals to identify a large catalogue of SNPs: WGS of 243 chickens from 24 chicken lines derived from diverse sources lead to the detection of about 139 million putative SNPs (Kranis et al., 2013).

In this study, the average distance between two successive variants was 60 bp, indicating important polymorphism level in the rainbow trout genome. This is consistent with the average SNP rate over all chromosomes of one SNP every 64 bp previously reported by Gao et al. (2018) in the Swanson rainbow trout reference genome. Such short average distance between successive variants was a strong limiting factor to preselect SNPs to design the HD chip. Indeed, an important technical issue in SNP array design is that very high SNP densities can potentially cause allele dropout when genotyping due to interferences between polymorphism at the marker position and at the probe designs that have to be monomorphic sequences flanking the marker candidates. When searching for markers with intermarker distance over 50 bp that could be considered in the HD array design, we could only retain 3.68 M SNPs.

Across all chromosomes, the average SNP density per Mb was 16,483 SNPs, i.e. slightly higher than the ~15,600 SNPs per Mb reported by Gao et al. (2018), although density of SNPs varied strongly from one chromosome to another (from 13,200 for Omy26 to 20,132 for Omy22). Interestingly, the lower SNP densities on Omy26 was also described in Gao et al. (2018) and associated with a higher proportion of SNPs being filtered out as putative paralogous sequence variants (PSV), as this chromosome shares high sequence homology with other chromosome arms in the genome as a result of delayed re-diploidization. Stronger variation in average SNPs density among chromosomes has been reported previously in chickens (Kranis et al., 2013) and humans (Zhao et al., 2003), with average value of 78 and 83.3 SNPs per kb across the genome but with some chromosomes having only 3 (on chromosome Z) and 2 (on chromosome Y) SNPs reported per kb respectively.

There was also a heterogeneous distribution of SNPs along the chromosomes, with a minimum density per Mb as low as 451 SNPs, and a maximum of 31,819 SNPs. Areas with less SNP density generally located at the telomeric parts or the centromeric parts (for metacentric chromosomes) of some chromosomes (e.g. Omy13 and Omy14) (Figure 3). Such heterogeneous distribution of SNPs has been previously reported in Eukaryotes, with potential explanations including heterogeneous recombination across the genome. It has been reported in a meta-analysis in eukaryotes that “heterogeneity in the distribution of crossover across the genome is a key determinant of heterogeneity in the distribution of genetic variation within and between populations” (Haenel et al., 2018). One broad-scale and general pattern observed within chromosomes is a lower recombination rate around centromeres (Stapley et al., 2017) and higher rates at the telomeric parts (Sakamoto et al., 2000; Anderson et al., 2012). Because higher recombination rates are observed in telomeric than centromeric regions of chromosomes, a higher number of variants may be expected in the telomeres. However, in general, the telomeres have very long patterns of repeats which generate problems in reads mapping. In the centromeric regions, it is unclear whether or not suppressed recombination is linked to highly repetitive regions (Talbert and Henikoff, 2010). Last but not least, the complexity of the rainbow trout genome with its recent whole genome duplication and partial rediploidization, and patterns of tetrasomic inheritance (Pearse et al., 2019), can potentially explain the difficulties to sequence and assemble some parts of its genome and hence detect SNPs. In a recent paper, Gui et al. (2022) have reported several phenomena (such as massive sequence divergences, extensive chromosome rearrangements, large-scale transposon bursts) occurring during the polyploidization and rediploidization that could explain the difficulties in assembling the complex genomes of Salmonids and other tetraploid fish species. Indeed, rainbow trout has a high content (57.1%) of repetitive sequences (Pearse et al., 2019), similar to the 59.9% reported for Atlantic salmon (Lien et al., 2016).

Taking advantage of the biological characteristics of fish (external fertilization and embryonic development, viability of uniparental progeny), isogenic lines have been generated in some fish species (reviewed in Franěk et al., 2020), by either gynogenesis (Quillet et al., 2007) or androgenesis (Young et al., 1996) in rainbow trout. Both USDA and INRAE datasets included the sequencing of 11 and 19 doubled-haploid individuals respectively from 30 different isogenic lines. This number, quite large and unique in fish, makes it possible to take advantage of both the within-line characteristics (homozygosity, isogenicity) and between-line variability. In particular, rainbow trout isogenic lines are being used for the development of genomic tools: the trout genome is the result of a whole genome duplication event that occurred about 96 Mya ago (Berthelot et al., 2014). Therefore, many genomic regions remain in a pseudo-tetraploid status, which complicates sequence assembly and development of genetic markers because of the difficulty to distinguish true allelic variants from PSVs. Therefore, homozygous individuals were used to produce the first genome sequence and reference transcriptome (Berthelot et al., 2014), subsequent improved genome assemblies (Pearse et al., 2019; Gao et al., 2021), and also to validate the large set of SNPs used in the first 57K SNP chip (Palti et al., 2014, 2015). In the present study, as in Gao et al. (2018), putative PSVs were filtered out by using genotypes informations from the isogenic lines, in order to generate a comprehensive catalogue of reliable SNPs in rainbow trout and then filter out SNPs to be included onto the HD SNP chip.

The 665K SNP chip was designed based on the Swanson reference genome (Pearse et al., 2019). Only 576K SNPs were uniquely positioned on the Arlee reference genome, which led to a few gaps over 1 Mb based on this reference genome (Figure 5) while there was no gap over 250 kb on the Swanson reference genome (Figure 2). Genetic and genomic differences between the Swanson and Arlee lines have previously been studied (Palti et al., 2014). It is also known that the two lines differ in their chromosomes’ numbers, the Swanson line having 2N=58 with 29 haploid chromosomes (Phillips and Ráb, 2001) and the Arlee line 2N=64 with 32 haploid chromosomes (Ristow et al., 1998). This is not surprising as there are some variable chromosome numbers in rainbow trout populations, associated with Robertsonian centric fusions or fissions, as for instance fission splitting metacentric chromosome 25 observed in Swanson genome into two acrocentric chromosomes in French lines (Guyomard et al., 2012; D’Ambrosio et al., 2019). Depending on the rainbow trout populations, the number of haploid chromosomes (N) varies from 29 to 32 and evidence suggests that the redband trout with 2N =58 is the most ancestral type (Thorgaard et al., 1983). In the Arlee karyotype the haploid chromosome number is 32 because chromosomes Omy4, Omy14 and Omy25 are divided into six acrocentric chromosomes (Gao et al., 2021). Note that Arlee chromosomes Omy30, Omy31 and Omy32 correspond to the p-arms of, respectively, Omy4, Omy25 and Omy14 on the Swanson genome.

The 664,531 SNPs successfully genotyped on 463 individuals across two French commercial populations represent a valuable tool for ongoing genomic studies on the genomic architecture of traits, the population evolution history and genetic diversity as well as for the assessment of inbreeding and the genetic effects of management practices in farmed populations. The HD chip is a powerful genomic tool that allow not only to have on average all along the genome a very high density of markers in comparison to the 57K chip, but also to significantly reduce the number of large gaps (> 1 Mb) in the genome coverage. In particular, the extremely low coverage at the telomeric parts of most of the chromosomes or at the centromeric part of metacentric chromosomes have been drastically reduced and the 2 regions spanning over 10 Mb each without any markers on Omy13 (see Supplementary Figure 1) have been drastically reduced, leaving just a large gap of 2.75 Mb at the end of Omy13 on the Arlee reference genome. This remaining gap is likely due to the fact that the entire chromosome Omy13 shares high sequence homology with other chromosome arms due to delay in re-diploidization (Gao et al., 2018). The next step will be to develop a new medium-density SNP array for rainbow trout keeping the 39K SNPs present on both the HD chip and the initial 57K chip, but adding about 25K SNPs of the HD chip to fill the large gaps without any SNP of the 57K chip. This second version of the medium-density chip will be a very useful tool both for genomic selection and for cost-effective GWAS thanks to imputation to HD genotypes.

In our study, we illustrate the interest of the HD chip based on LD study across three different rainbow trout populations. The analysis of LD plays a central role in GWAS and fine mapping of QTLs as well as in population genetics to build genetic maps, to estimate recombination rates or effective population sizes as the expected value of *r*^2^ is a function of the parameter 4N_e_c, where *c* is the recombination rate in Morgan between the markers and *N*_e_ is the effective population size (Sved, 1971). The decay and extent of LD at a pairwise distance can be used to determine the evolutionary history of populations (Hayes et al., 2003; Santiago et al., 2020). Lines LB and LC had the highest LD values in comparison to the American hatchery population (HA), potentially indicating lower effective population sizes in the French selected lines. The lower LD values in the American population may be partly linked to stratification in the sampled population gathered from diverse rivers, but however it helps to quantify the lower bound LD values at short distance that we may expect in hatchery populations. The higher than average LD observed on Omy5 is likely caused by a large chromosomal double-inversion of 55 Mb (Pearse et al., 2019) which prevents recombination in fish. While a number of studies quantify in salmonids the presence of long-range LD from 50 kb to over 1 Mb either for commercial populations (Kijas et al., 2017; Vallejo et al., 2018; Barría et al., 2019; D’Ambrosio et al., 2019) or wild populations (Kijas et al., 2017), little is known on the LD at very short distances. In rainbow trout farmed populations, the level of strong LD (*r*^2^ > 0.20) spans over 100 kb (D’Ambrosio et al., 2019) to 1 Mb (Vallejo et al., 2018, 2020).

Barría et al. (2019) indicated a maximum value of 0.21 in a Chilean Coho selected line for marker distance lower than 1 kb and a threshold value of r2=0.2 reached at approximately 40 kb. In Atlantic salmon, r2=0.2 was reached at approximately 200 kb in a Tasmanian farmed population coming from a single Canadian river without any further introgression (Kijas et al., 2017). In the Tasmanian population, the average LD value for markers separated by 0–10 kb was 0.54 while the corresponding average LD value was only 0.04 in a Finish wild population (Kijas et al., 2017).

In our study, regardless of the rainbow trout populations, the LD values at very short distances between markers (≤ 10 kb) were moderate (0.44 - 0.47 at 2 kb and 0.34-0.38 at 10 kb, respectively for LB and LC) compared to the ones observed at similar distances in cattle breeds (Hozé et al., 2013) where r2 values were around 0.70 at 2 kb and in the range 0.50-0.55 at 10 kb whatever the breeds considered. This may be partly due to higher recombination rate in rainbow trout (1.67 cM/Mb; D’Ambrosio et al., 2019) than in cattle (1.25 cM/Mb; Arias et al., 2009), but it also indicates that the founder populations of rainbow trout farmed lines have presumably larger ancestral effective population sizes than cattle breeds. On the contrary, for marker distances over 100 kb, LD values decrease below 0.20 in cattle breeds, while average LD values are still 0.26 to 0.30 in LC and LB lines, respectively. This indicates stronger recent bottlenecks and selection rates in rainbow trout lines than in cattle breeds. Similar long-range LD was independently observed in two US commercial rainbow trout populations (Vallejo et al., 2018, 2020). The pattern of LD decay in rainbow trout commercial lines appears to be more similar to the one observed in conservation flocks of chicken from South Africa (Khanyile et al., 2015), with very similar values reported both at shorter distances than 10 kb, as well as at 500 kb distance where LD values range from 0.15 to 0.24 depending on the conservation flocks and values of 0.21 to 0.25 were derived for LC and LB, respectively. A last factor that may contribute to this long-range LD in rainbow trout is the high crossing-over interference in males observed when plotting the linkage map distance between markers from the male vs. female linkage maps against the physical distance in base pairs. Sakamoto et al. (2000) have reported a 3.25:1 female to male linkage map distance ratio and Gonzalez-Pena et al. (2016) indicates that female/male recombination ratios were above 2.0 in all the 13 chromosomes known to have homologous pairing with at least one other chromosome arm, while in most of the non-duplicated chromosomes the ratio was generally lower. Because such high crossing-over interference in males were observed in families generated from sex-reversed XX males, we hypothesize that there must be a mechanism that is controlling meiosis in the sperm differently than in the eggs through a different regulation of gene expression not related to presence or absence of the sdY gene.

We have demonstrated in this paper a substantial linkage disequilibrium between neighboring markers, suggesting the density of genotyped SNPs is well-designed to accurately tag most areas of the rainbow trout genome. We acknowledge that, by design, the minor allele frequency distribution of genotyped SNPs is skewed to common alleles, and variation has been predominantly sampled from common SNP shared by both French and North American farmed populations. While this may limit some analyses, we believe that the array will be an invaluable genomic resource for ongoing work investigating genetic diversity, genetic architecture of traits and adaptive potential in world-wide rainbow trout populations.

## 6 Author Contributions

M.B., D.L. and F.P. conceived and designed the study. M.D.N, E.Q. and P.H. were involved in the conceptualisation and funding acquisition for the project. Y.P. and G.G. gave access to the USDA SNP databases. M.B., A.D., M.C. and D.L. performed bioinformatics analyses on resequencing data. M.B., A.D., G.G. and F.P led the design of the 665K SNP array. P.P. coordinated the collection of samples to be genotyped with the 665K SNP array. J.D., L.J. and C.P. performed the 665K SNP genotyping. K.P., H.L., M.P. and F.P. analyzed the 665K genotyping data. M.B., D.L. and F.P. wrote the manuscript. All authors reviewed and approved the manuscript.

## 8 Acknowledgments

The SNP chip was developed in cooperation with Thermo Fisher and we particularly thank the following Thermo Fisher Scientific personnel for their direct contribution: Ruth Barral Arca, Marie-Laure Schneider and Philippe Lavisse. We are also grateful to the Genotoul bioinformatics platform (Toulouse Occitanie, doi:10.15454/1.5572369328961167E12) and the INRAE MIGALE bioinformatics facility (MIGALE, INRAE, 2020. Migale bioinformatics Facility, doi: 10.15454/1.5572390655343293E12) for providing help, computing and storage resources. We also thank the 3 French breeding companies “Les fils de Charles Murgat”, “Bretagne Truite” and “Viviers de Sarrance” that provided samples for genome resequencing or genotyping on the 665K SNP array.

## 10 Data Availability Statement

Raw sequence data that were generated for French isogenic lines are deposited in the ENA (Projects PRJEB52016 and PRJEB51847).

The VCF file for the database of all the SNPs identified in this study including a file with allele frequency information for each SNP in the database are available for downloading from a public repository (dataINRAE).

The sequence and the genotypes of the three French commercial trout lines from “Les Fils de Charles Murgat” (Beaurepaire, France), “Bretagne Truite” (Plouigneau, France) and “Vivers de Sarrance” (Sarrance, France) breeding companies will be made available by request on the recommendation of Pierrick Haffray (SYSAAF).

## Supplementary Material

**Supplementary Figure 1.**
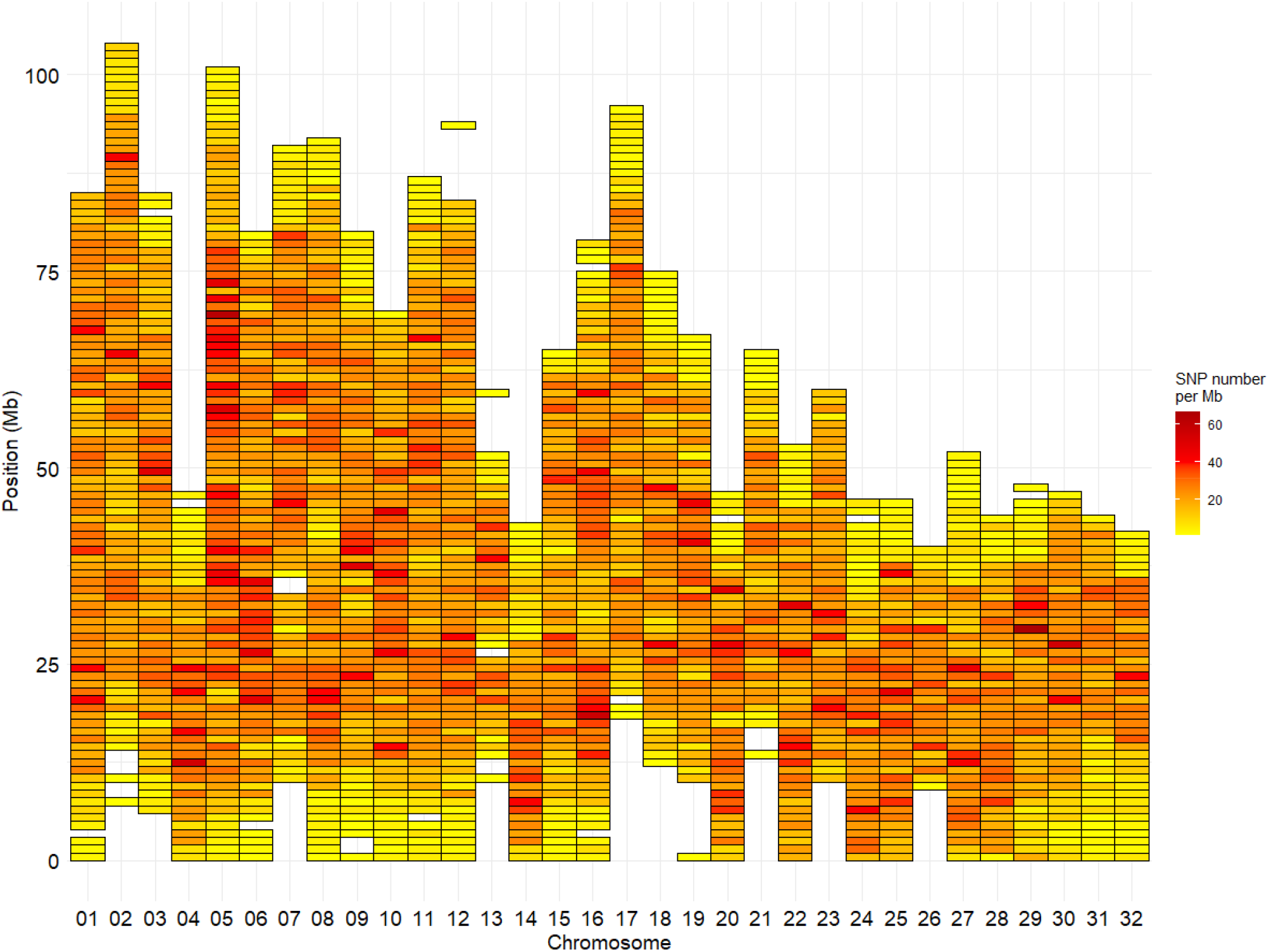
Marker density per Mb for the LD Trout Affymetrix array with 38,948 SNPs positioned on the 32 chromosomes of the Arlee genome reference

